# Mistranslating tRNA variants impact the proteome and phosphoproteome of *Saccharomyces cerevisiae*

**DOI:** 10.1101/2025.09.15.676438

**Authors:** Matthew D. Berg, Alexis T. Chang, Ricard A. Rodriguez-Mias, Judit Villén

## Abstract

Transfer RNAs (tRNAs) ensure accurate decoding of the genetic code. However, mutations in tRNAs can lead to mis-incorporation of an amino acid that differs from the genetic message in a process known as mistranslation. As mistranslating tRNAs modify how the genetic message is decoded, they have potential as therapeutic tools for diseases caused by nonsense and missense mutations. Despite this, they also produce proteome-wide mis-made proteins which can disrupt proteostasis. To better understand the impact of mistranslating tRNA variants, we profile the proteome and phosphoproteome of yeast expressing three different mistranslating tRNAs. While the overall impacts were similar, the extent of growth defects and proteome changes varied with the substitution type. Although the global impacts were modest, mistranslation influenced key cellular processes, including proteostasis, cell cycle and translation. These findings highlight the need to consider cellular consequences when developing mistranslating tRNAs for therapeutic applications.

## INTRODUCTION

Transfer RNAs (tRNAs) maintain the fidelity of the genetic code, ensuring that the amino acid specified by the mRNA is incorporated into the growing polypeptide chain (reviewed in Berg and Brandl 2021). Mutations in tRNAs can lead to mistranslation, the mis-incorporation of an amino acid that differs from what is specified by the genetic message. Early studies identified mistranslating tRNAs as suppressors of nonsense and missense mutations (Stadler and Yanofsky 1959; Yanofsky and Crawford 1959; Crawford and Yanofsky 1959; Benzer and Champe 1962; Gorini and Beckwith 1966). For example, Goodman *et al*. (1977) found an anticodon mutation in yeast tRNA^Tyr^ that suppresses ochre stop codons and others discovered *Escherichia coli* tRNA variants that insert glycine at cysteine or arginine codons (Carbon *et al*. 1966; Gupta and Khorana 1966).

Because mistranslating tRNAs alter the reading of the genetic message, they are currently being developed as therapeutics for diseases caused by nonsense and missense mutations (reviewed in Coller and Ignatova 2024). Temple et al. (1982) first demonstrated that a mutant tRNA^Lys^ which suppresses amber stop codons could treat β-thalassemia. More recently, several groups engineered tRNAs to suppress premature termination codons that cause a variety of diseases including cystic fibrosis (Lueck *et al*. 2019; Ko *et al*. 2022; Albers *et al*. 2023), Ullrich disease (Sako *et al*. 2006), hereditary diffuse gastric cancer (Bordeira-Carriço *et al*. 2014), human mucopolysaccharidosis type I (Wang *et al*. 2022), hereditary breast and ovarian cancer syndrome (Awawdeh *et al*. 2024; Specht *et* al. 2025), and frontotemporal dementia (Beharry *et al*. 2025). Furthermore, to treat diseases caused by missense mutations, Hou *et al*. (2023) developed a variety of missense-correcting tRNAs including variants capable of suppressing mutations associated with limb-girdle muscular dystrophy type 2A.

In addition to their therapeutic applications, mistranslating tRNAs naturally exist as both rare and common variants in human populations and could contribute to disease (Berg *et al*. 2019a; Hasan *et* al. 2023; Tennakoon et al. 2025). While the effects of mistranslation tRNA variants are often buffered by multiple tRNA gene copies, mistranslation disrupts proteostasis and induces harmful impacts in model systems including slow growth in yeast (Paredes *et al*. 2012; Hoffman *et al*. 2017; Berg *et al*. 2017, Berg *et al*. 2021a; McDonald *et al*. 2025), deformities and decreased viability in flies and zebrafish (Reverendo *et al*. 2014; Isaacson *et al*. 2022, Isaacson *et al*. 2024), and cardiac abnormalities and neurodegeneration in mice (Lu *et al*. 2014; Liu *et al*. 2014). We have also found that different genetic backgrounds can exacerbate the impacts of mistranslating tRNA variants (Berg *et al*. 2021b, Berg *et al*. 2022).

Despite the prevalence of mistranslating tRNA variants in human populations and their growing popularity as therapeutics, their impact on cellular physiology is poorly defined particularly at the proteome level. In this report, we investigate the proteomic and phosphoproteomic response to different mistranslating tRNA variants. Using three mistranslating tRNAs expressed in *Saccharomyces cerevisiae* that each create different substitutions, we find that the number of protein and phosphosite abundance changes correlate with the growth impact of each tRNA variant. In addition to upregulating heat shock proteins, expressing mistranslating tRNA variants impacts abundance and phosphorylation of proteins involved in cell cycle and translation. These results will guide future studies into mechanisms through which tRNA variants contribute to disease and inform potential side effects of tRNA therapeutics.

## MATERIALS AND METHODS

### Yeast strain and growth

Wild type haploid yeast strain BY4741 (*MATa his3Δ1 leu2Δ0 met15Δ0 ura3Δ0*; Brachmann *et al*. 1998) is a derivative of S288c. Strains containing mistranslating tRNAs were made by transforming BY4741 with *URA3* plasmids encoding tRNA^Pro^_G3:U70_ (pCB2948; Hoffman *et al*. 2017), tRNA^Ser^ _UGG,G26A_ (pCB4023; Berg *et al*. 2017) or tRNA^Ser^ _UCU,G26A_ (pCB4257; Berg *et al*. 2019b). An empty vector (YCplac33; Gietz and Sugino 1988) was transformed into BY4741 to create the control strain.

Yeast strains were grown at 30°C in yeast peptone medium containing 2% glucose or synthetic media supplemented with nitrogenous bases and amino acids unless otherwise indicated. For growth curves, cells were grown to stationary phase, diluted to OD_600_ ∼ 0.1 in minimal media and incubated at 30°C. Every 15 minutes, OD_600_ was measured using a BioTek Epoch 2 microplate spectrophotometer for 24 hours. Doubling time was calculated using the R package ‘growthcurver’ (Sprouffske and Wagner 2016).

### Whole proteome and phosphoproteome sample preparation

Six replicates of each strain were grown in synthetic minimal media lacking uracil. Overnight cultures were diluted to OD_600_ ∼ 0.1 and harvested at an OD_600_ between 0.8 and 1.0 by adding 100% (w/v) trichloroacetic acid directly to the liquid culture to a final concentration of 10%. Cultures were incubated on ice for 10 minutes, centrifuged, washed once with ice cold acetone, centrifuged again, and washed with ice cold water. Cell pellets were snap frozen in liquid nitrogen.

Cell pellets were resuspended in a denaturing lysis buffer (8 M urea, 50 mM Tris pH 8.2, 75 mM NaCl). Cells were lysed with 0.5 mm zirconia/silica beads for four 1-minute cycles of bead beating with 1 minute rest on ice in between. Lysate was cleared by centrifugation at 21,000 x g for 10 minutes at 4°C. Protein concentration was determined by bicinchoninic acid assay (BCA; Pierce, ThermoFisher Scientific). Proteins were reduced with 5 mM dithiothreitol for 30 minutes at 55°C, alkylated with 15 mM iodoacetamide for 30 minutes at room temperature in the dark, and quenched with an additional 5 mM dithiothreitol for 30 minutes at room temperature.

Lysates were processed with the R2-P1 protocol (Leutert *et al*. 2019) implemented on a KingFisher Flex (ThermoFisher) magnetic particle processing robot with minor modifications. Briefly, lysates were block randomized across a 96-deep well plate and 300 μg protein was combined with 600 μg carboxylated paramagnetic beads (1:1 mix of hydrophilic and hydrophobic Sera-Mag SpeedBead Carboxylated-Modified, GE Life Sciences) and ethanol was added to 80%. Proteins were washed four times in 80% ethanol before digestion in 50 mM triethylammonium bicarbonate pH 8.5 for 14 hours at 37°C. Samples prepared to assess mistranslation frequency were digested with 5 ng/μL LysC (Wako Chemicals) and samples prepared for whole proteome and phosphoproteome analysis were digested with a combination of 5 ng/μL trypsin (Promega) with 5 ng/μL LysC. Digests were acidified to pH ∼ 2 with formic acid (FA), 5% of the sample was taken out for total proteome analysis and desalted over Empore C18 stage tips (Rappsilber et al. 2007). All samples were dried down.

To enrich phosphopeptides, 275 μg of dried trypsin/LysC digested peptides were resuspended in 80% acetonitrile (ACN) with 0.1% trifluoroacetic acid (TFA). Following clarification by centrifugation, R2-P2 (Leutert *et al*. 2019) was performed on a KingFisher Flex. Briefly, phosphopeptides were enriched with Fe^3+^-NTA magnetic beads (PureCube Fe-NTA MagBeads, Cube Biotech), washed three times with 80% ACN, 0.1% TFA and eluted in 50% ACN, 2.5% NH_4_OH. Enriched phosphopeptides were acidified and dried down.

### Mass spectrometry data acquisition

Peptides were resuspended in 4% ACN, 3% FA and analyzed on an Orbitrap Eclipse Mass Spectrometer (ThermoFisher Scientific) equipped with an Easy1200 nanoLC system (ThermoFisher Scientific). Peptides were loaded onto a 100 μm ID x 3 cm precolumn packed with Reprosil C18 3 μm beads (Dr. Maisch GmbH) and separated by reverse-phase chromatography on a 100 μm ID x 30 cm analytical column packed with Reprosil C18 1.9 μm beads (Dr. Maisch GmbH) housed in a column heater set at 50°C using two buffers: (A) 0.1% FA in water and (B) 80% ACN in 0.1% FA in water at 450 nL/min flow rate.

For whole proteome measurements, peptides were separated over a 120 min gradient ramping from 7% to 50% B over 103 min and washing at 90% B before re-equilibrating to 3%. For data independent acquisition (DIA) measurements, a staggered window approach was used (Amodei *et al*. 2019). A full MS1 scan was recorded after every DIA cycle at 60,000 resolution with standard automatic gain control (AGC) target and automatic injection time (IT). Two distinct offset DIA cycles shifted 12 m/z covered an effective range of 363 to 1095 m/z. One DIA cycle contained 30 windows of 24 m/z size at 30,000 resolution, 33% normalized collision energy (NCE) higher-energy collisional dissociation (HCD), default charge state 3 and AGC target of 1000%. For gas phase fractionated (GPF) samples, eight injections of a pooled sample covering 100 m/z of the total scan range per injection with 25 windows of 4 m/z isolation width were performed. For data dependent acquisition (DDA) measurements, a full MS1 scan was recorded every 3 sec at 120,000 resolution with standard AGC target and automatic IT from 375 to 1500 m/z. MS2 scans were recorded at 30,000 resolution, 30% NCE HCD, 1.6 m/z isolation window, standard AGC target and automatic IT.

For phosphoproteome measurements, phosphopeptides were separated over a 90 min gradient ramping from 6% to 50% B over 73 min and washing at 90% B before re-equilibrating to 3%. For DIA measurements, a staggered window approach was used with two distinct offset DIA cycles shifted 12 m/z covering an effective range of 438 to 1170 m/z. One DIA cycle contained 30 windows of 24 m/z size at 50,000 resolution, 31% NCE HCD, default charge state 3 and AGC target of 1000%. A full MS1 scan was recorded after every DIA cycle at 60,000 resolution with standard AGC target and automatic IT. For DDA measurements, a full MS1 scan was recorded every 3 sec at 120,000 resolution with standard AGC target and automatic IT from 375 to 1500 m/z. MS2 scans were recorded at 50,000 resolution, 30% NCE HCD, 1.6 m/z isolation window, standard AGC target and automatic IT.

### Mass spectrometry data analysis

MS raw files were converted to mzML using MSconvert v3.0.23305.fafbb32 (Adusumilli and Mallick 2017). Staggered DIA files were converted using filters peakPicking at vendor msLevel 1- and demultiplexed with optimization overlap_only and massError 10 ppm. Database and spectral library search were performed using Fragpipe v22.0 (Yu *et al*. 2023) with *S. cerevisiae* FASTA file downloaded from Uniprot on 2024-04-22. To quantify mistranslation, DDA files were searched with default settings for LysC and high resolution MS data with variable modifications of methionine oxidation (15.9949 Da), protein N-terminal acetylation (42.0106 Da), proline to alanine (-26.0156 Da), proline to serine (-10.0207 Da) and arginine to serine (-69.0690 Da). To detect new phosphosites, DDA files from phospho-enriched proteome samples were searched with default settings for trypsin and high resolution MS data with variable modifications of serine, threonine and tyrosine phosphorylation (79.9663 Da), proline to alanine (-26.0156 Da), proline to serine (-10.0207 Da), arginine to serine (-69.0690 Da), proline to phosphoserine (69.9456 Da) and arginine to phosphoserine (10.8972 Da). PTM site localization was enabled using PTMProphet for all modifications. For whole proteome samples, DIA, DDA and DIA-GPF files were used to create a spectral library in Fragpipe. Default settings for trypsin and high resolution MS data were used with variable modifications of methionine oxidation (15.9949 Da) and protein N-terminal acetylation (42.0106 Da). DIA files were searched and quantified with DIA-NN v2.1.0 (Demichev *et al*. 2020) using the spectral library created in FragPipe in “QuantUMS (high precision)” mode. For phosphoproteome samples, DIA and DDA files were combined with raw files from the deep yeast phosphoproteome samples generated by Leutert *et al*. (2023) to create a spectral library in Fragpipe as above but including variable modification of phosphorylation on STY residues (79.9663 Da).

Data analysis was performed using custom scripts in RStudio v2023.03.1 and R v4.3.0. Summary plots were made with modified code from the ‘protti’ package (Quast *et al*. 2022). Mistranslation frequency was calculated from the whole proteome DDA data using unique mistranslated peptides for which the non-mistranslated sibling peptide was also observed. Mistranslation frequency is defined as the counts of unique mistranslated peptides divided by the counts of all peptides containing the target amino acid or codon and expressed as a percentage. The whole proteome DIA data was filtered at ≤ 1% FDR with Global.Q.Value at the precursor level. For the phosphoproteome DIA data, precursors were filtered at ≤ 1% FDR with Global.Q.Value and Peptidoform.Q.Value and only sites with PTM.Site.Confidence greater than 0.75 were retained. MSstatsPTM (Kohler *et al*. 2022) was used to impute missing values, summarize up to protein and phosphopeptide levels and to determine differentially abundant proteins and phosphopeptides. Gene ontology (GO) and wikipathway enrichment was performed using g:profiler (Kolberg *et al*. 2023) with all quantified proteins used as the background.

Report logs containing full parameters for the raw data analysis, DIA libraries, the DIA-NN outputs and custom R scripts used to analyze the data and create the figures can be found at https://github.com/Villen-Lab/Mistranslation-Phospho-Proteome-AnalysisFiles.

## RESULTS AND DISCUSSION

### Frequency of mistranslation does not correlate with growth impact of tRNA variants

To understand the impact of mistranslating tRNAs and the pathways cells induce to cope with the loss of translation fidelity, we used mass spectrometry to measure the proteome and phosphoproteome of three yeast strains expressing different mistranslating tRNA variants. The tRNA variants are shown in Figure 1A. The first is a proline tRNA with a G3:U70 base pair in its acceptor stem that mis-incorporates alanine at proline codons (Pro→Ala; Hoffman *et al*. 2017). The other two are serine tRNAs with either a UGG proline or UCU arginine anticodon which mis-incorporate serine at proline (Pro→Ser) or serine at arginine codons (Arg→Ser), respectively (Berg *et al*. 2017, 2019b). The serine tRNA variants also have a G26A mutation which prevents a key nucleotide modification, dampening tRNA function and allowing tolerable mistranslation levels (Berg *et al*. 2017).

**Figure 1.**
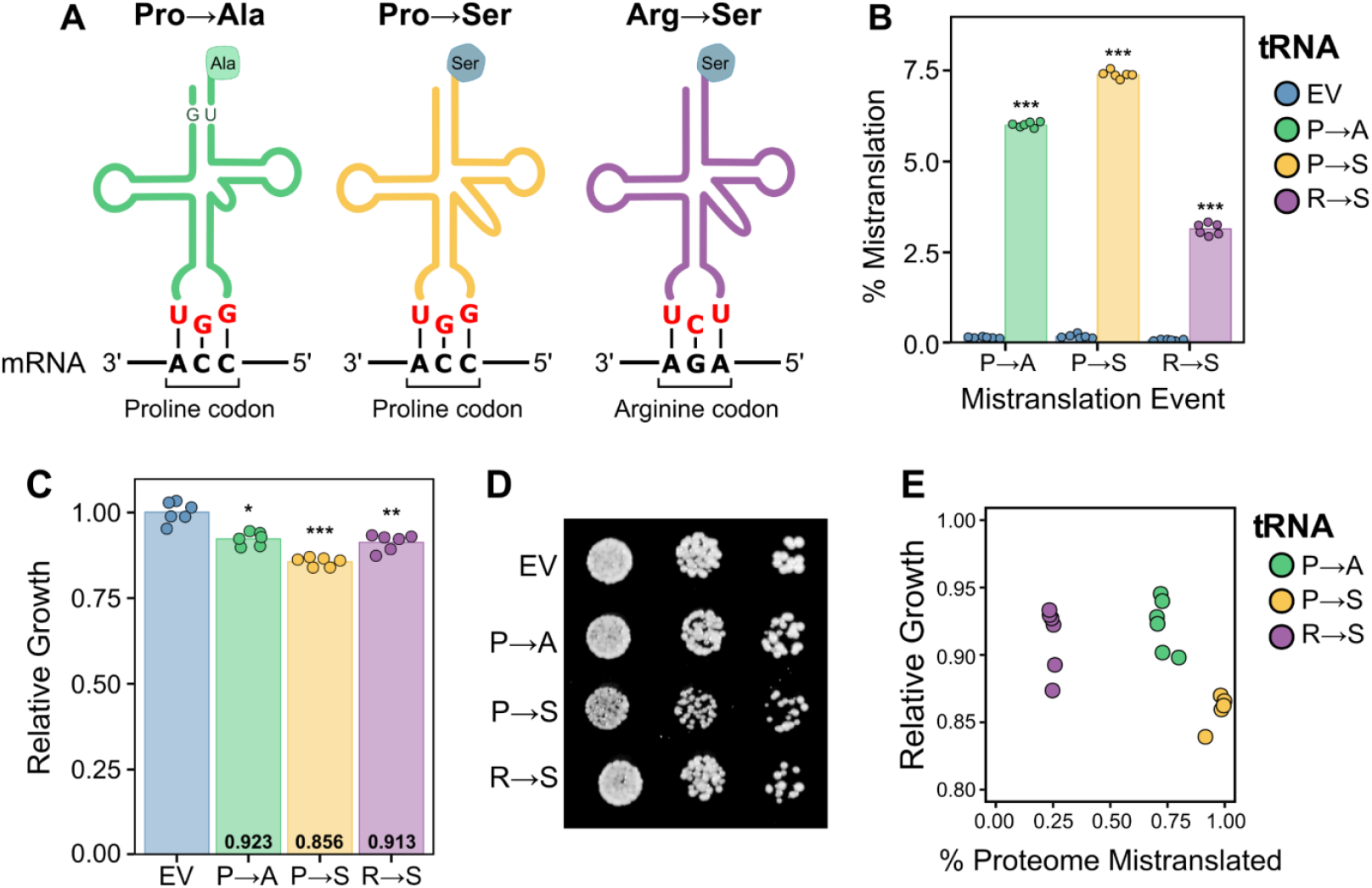
tRNA variants induce mistranslation. **(A)** Schematic representing the three mistranslating tRNAs used in this study. **(B)** Mistranslation frequency for each strain determined by mass spectrometry. Wild-type yeast strain BY4741 expressing either an empty vector control or a mistranslating tRNA variant were grown in minimal media lacking uracil to an OD_600_ of ∼ 1.0 and mass spectrometry analysis of the cellular proteome was performed. The percent of unique peptides detected with either proline to alanine, proline to serine or arginine to serine substitutions was calculated for each respective variant and the control strain. **(C)** Growth rate for the control strain and strains expressing each mistranslating tRNA was determined from growth curves performed in minimal media lacking uracil. Strains were diluted to an OD_600_ of ∼ 0.1 and grown for 24 hours. Growth relative to the control strain, set as 1.0, was calculated from the doubling time. **(D)** Representative growth of the control strain and strains expressing each mistranslating tRNA on solid minimal media lacking uracil. Overnight cultures of each strain were grown in minimal media lacking uracil, spotted in 10-fold serial dilutions and grown for two days. **(E)** Correlation between relative growth rate of each strain as determined in (C) versus mistranslation frequency calculated by summing the total intensity of all identified mistranslated peptides and normalized to the total intensity of peptides identified in the mass spectrometry data. In all panels, each point represents one biological replicate. Strains expressing a mistranslating tRNA were compared to the empty vector (EV) control strain using a *t-*test with Bonferroni correction. * *P* < 0.01; ** *P* < 0.001; *** *P* < 0.0001.

Using mass spectrometry, we first confirmed that all three strains had elevated levels of the expected substitution compared to a control strain containing an empty vector (Figure 1B and Table S1). Mistranslated peptides were found at a frequency of 6.0% for the Pro→Ala strain, 7.4% for the Pro→Ser strain, and 3.1% for the Arg→Ser strain based on unique counts of identified peptides. We also analyzed the subsets of codons being mistranslated in each strain (Figure S1 and Table S2). The two mistranslating tRNAs with UGG anticodons had similar profiles and, as we had previously observed for an alanine tRNA variant with proline UGG anticodon (Cozma *et al*. 2023), mistranslated at all four CCN proline codons with the majority of mis-incorporation occurring at the Watson-Crick complementary CCA codon. For the Arg→Ser tRNA variant, mis-incorporated serine was only observed at the Watson-Crick complementary AGA arginine codon.

As protein abundance and codon usage will influence the proportion of the proteome that is mistranslated, we summed the total intensity of all mistranslated peptides and normalized it to the total intensity of all identified peptides to estimate the proteome impact of each mistranslating tRNA. Mistranslated peptides comprised 0.73 ± 0.03% of the total measurable proteome for the Pro→Ala tRNA, 0.98 ± 0.03% for the Pro→Ser tRNA and 0.25 ± 0.01% for the Arg→Ser tRNA. We also determined the growth impact of each mistranslating tRNA. As shown in Figure 1C,D, the impact of these tRNAs on yeast growth was relatively minor. Interestingly, there was poor correlation between growth impact and amount of mistranslation (Figure 1E). Previously, we observed that mistranslation frequency correlates with growth impact when analyzing the same amino acid substitution type (Berg *et al*. 2019b). However, these results suggest that type of amino acid substitution impacts the cellular response to mistranslation consistent with what we and others have seen when analyzing tRNA variants that create different substitutions (Santos *et al*. 2018; Zimmerman *et al*. 2018; Cozma *et al*. 2023; Davey-Young et al. 2024; Isaacson *et al*. 2024).

### Characterizing the proteome and phosphoproteome of strains expressing mistranslating tRNA variants

To understand the proteome changes and phospho-signaling pathways induced in response to mistranslating tRNAs, we profiled the steady state cellular proteome and phosphoproteome of each mistranslating strain using a label free data independent acquisition (DIA) mass spectrometry approach (Figure 2A). On average, 4,661 ± 14 proteins and 10,290 ± 280 confidently localized phosphosites were quantified per sample (Figure 2B). The median coefficient of variation (CV) between six biological replicates was less than 15% for the proteome and less than 20% for the phosphoproteome (Figure 2C), indicating the measurements were precise and reproducible across samples. Principal component analysis conducted on the protein level measurements demonstrated that the mistranslating strains separated across PC1 with 25% of the variation explained in this dimension (Figure 2D). For the phosphoproteome measurements, the mistranslating strains separated across PC2 accounting for 16.8% of the variation. Interestingly, while the Pro→Ala and Arg→Ser samples clustered together in the proteome PCA, the Arg→Ser samples clustered separately from the Pro→Ala samples and closer to the empty vector control samples in the phosphoproteome PCA. Overall, these results indicate that our dataset captures distinct proteome and phosphoproteome changes that occur in response to mistranslating tRNAs.

**Figure 2.**
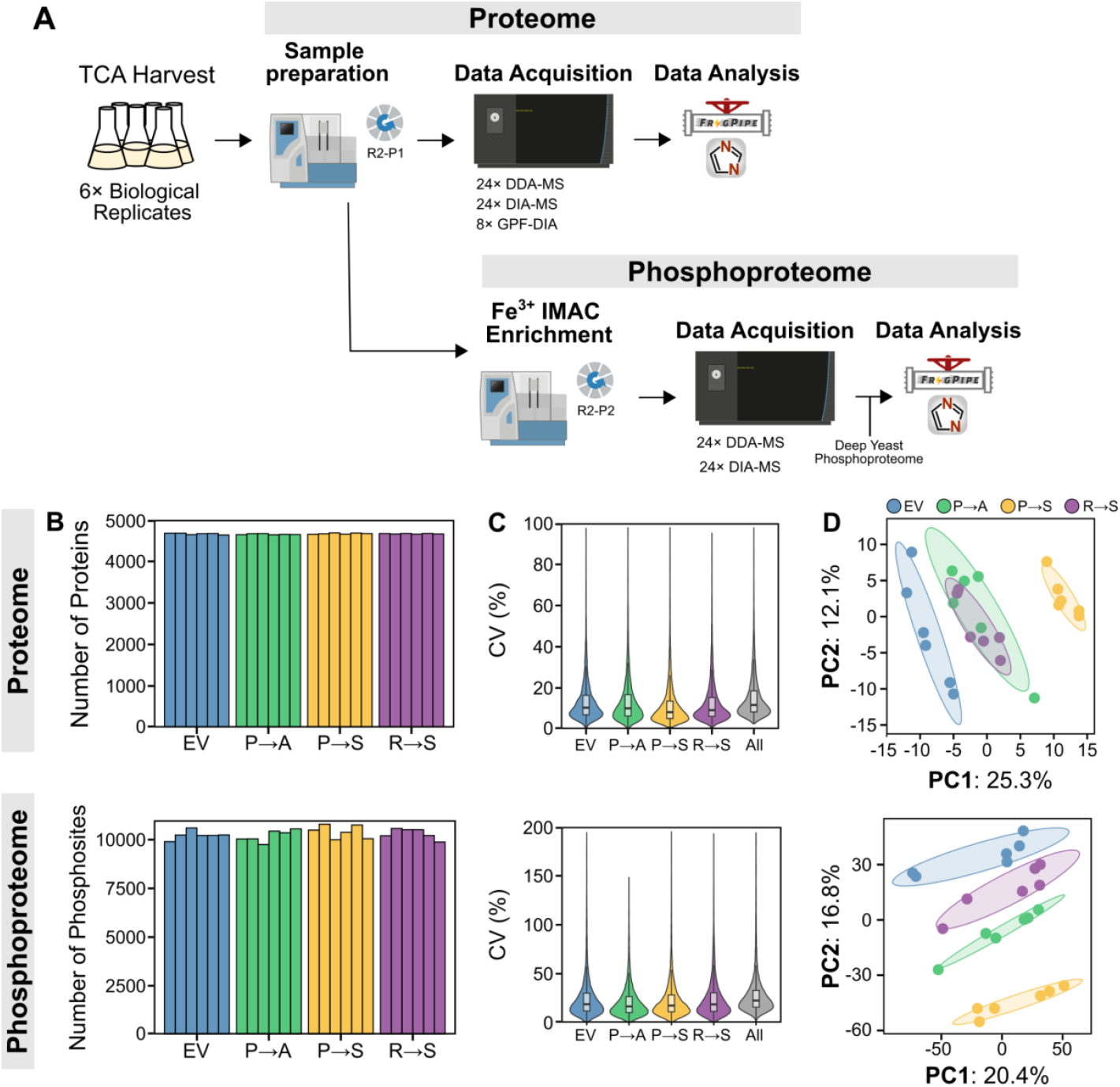
Sample preparation and dataset quality. **(A)** Workflow for analyzing proteome and phosphoproteome changes in response to mistranslation. Strains were grown in minimal media lacking uracil to an OD_600_ of ∼ 1.0 before cells were harvested with TCA. Proteins were digested into peptides and prepared for mass spectrometry using the R2-P2 protocol (Leutert *et al*. 2019). Proteome samples were measured by DDA and DIA. Eight gas phase fractions of a pooled sample were measured by narrow window DIA. Phosphoproteome samples were measured by DDA and DIA. All proteome data was used to create a proteome spectral library in Fragpipe (Yu *et al*. 2023). Phosphoproteome data was supplemented with deep yeast phosphoproteome samples from Leutert *et al*. (2023) to create a phosphoproteome spectral library. The DIA files were searched and quantified using the respective spectral libraries in DIA-NN (Demichev *et al*. 2020). **(B)** Total number of proteins (top) and phosphosites (bottom) quantified. Each bar represents one biological replicate. **(C)** Violin plot representing the distribution of coefficient of variation (CV) for all proteins (top) and phosphosites (bottom) quantified. **(D)** Principal component analysis of the whole proteome (top) and phosphoproteome (bottom) data from the empty vector control strain (EV) and strains expressing each mistranslating tRNA variant. Shaded ellipses indicate a one standard deviation confidence region for each group.

### Upregulation of proteostasis in response to mistranslating tRNAs

We first compared the proteomes of strains expressing each mistranslating tRNA to the empty vector control strain to identify differentially abundant proteins. Volcano plots highlighting significant protein abundance changes with adjusted *p*-values < 0.05 and absolute fold-changes > 1.5 are shown in Figure 3A. Protein abundances can be found in Table S3 and the fold-change and adjusted *p*-values for each protein in each condition can be found in Table S4. The Pro→Ser strain had the most differentially abundant proteins compared to control, with 142 proteins increased and 134 proteins decreased. Both the Pro→Ala and Arg→Ser strains had relatively few differentially abundant proteins, 65 and 58, respectively, consistent with their minimal impact on growth.

**Figure 3.**
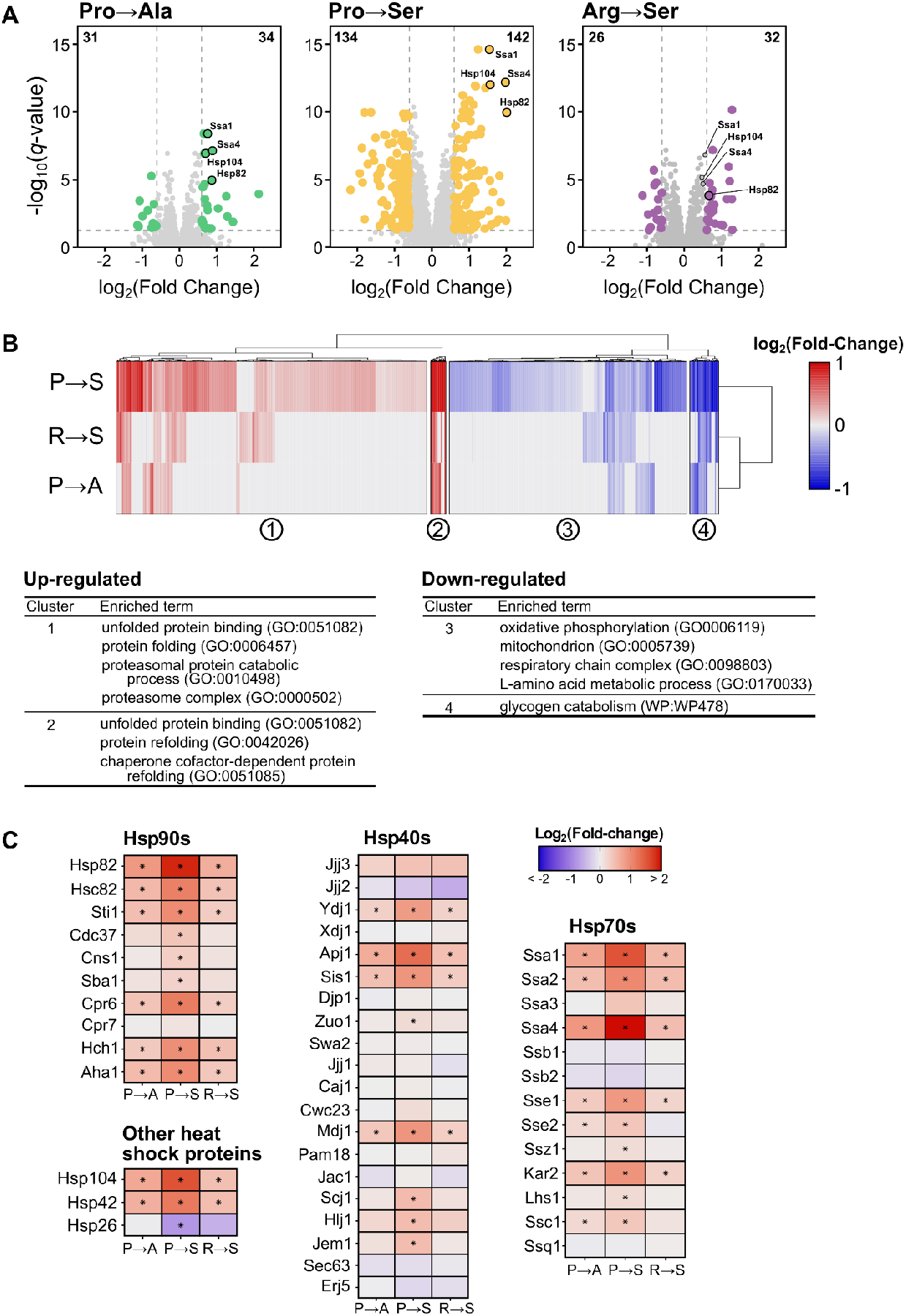
Proteome changes in response to mistranslating tRNA variants. **(A)** Volcano plots highlight differentially abundant proteins (adjusted *p-*value < 0.05, |log_2_ fold-change| > 0.5) for each mistranslating strain compared to the control. The most significantly up-regulated heat shock proteins are labeled. Numbers in the upper corners indicate the number of significantly up- and down-regulated proteins. **(B)** Heatmap of hierarchically clustered differentially abundant proteins (adjusted *p*-value < 0.05 in at least one strain) for each mistranslating strain relative to the control. Fold-change for each gene is the average of six replicates. Up-regulated proteins are colored red while down-regulated proteins are colored blue. Significantly enriched GO and wikipathway terms relative to all quantified proteins was determined for each cluster using g:Profiler (Kolberg *et al*. 2023). **(C)** Heat maps represent the log_2_-fold change in protein abundance for selected heat shock proteins. Stars indicate statistically significant changes in protein abundance relative to the empty vector control strain (adjusted *p-*value < 0.01).

To investigate similarities and differences in protein abundance changes globally amongst different mistranslating tRNA variants, proteins with significant abundance changes (adjusted *p*-value < 0.05) were hierarchically clustered based on their fold-change relative to the control strain and GO enrichment was performed on each cluster (Figure 3B). Overall, the proteome response to mistranslation is similar for different mistranslating tRNA variants with upregulation of protein quality control and downregulation of energy production.

We next looked closer at the up-regulated proteins involved in protein quality control. Specifically, the four most up-regulated chaperones in all strains were cytosolic Hsp70 chaperones Ssa1 (constitutively expressed) and Ssa4 (stress induced), the Hsp90 chaperone Hsp82 and the disaggregase Hsp104 (highlighted in Figure 3A). The magnitude of up-regulation of these proteins is proportional to the growth impact of each mistranslating tRNA with the Pro→Ser strain inducing the highest levels followed by the Pro→Ala strain. The Arg→Ser strain had relatively low induction. This trend also holds when looking more broadly at chaperone and co-chaperone proteins (Figure 3C). Only the small heat shock protein Hsp26, which binds unfolded proteins and is normally induced under stress conditions (Susek and Lindquist 1990), decreased in abundance in the Pro→Ser strain.

Looking more broadly at proteins involved in proteostasis, there was also a general increase in abundance for proteasome subunits in all strains relative to the control, supporting a role for protein turnover in dealing with mistranslated proteins (Figure S2A). In agreement with this, we previously identified negative synthetic genetic interactions between mistranslation and components of the proteasome (Hoffman et al. 2017; Berg *et al*. 2020, Berg *et al*. 2021b, Berg *et al*. 2022) and Kalapis *et al*. (2015) found cells adapt to mistranslation by increasing proteasome abundance. The largest increase in proteasome abundance was seen in the Pro→Ser strain, followed by the Arg→Ser and the Pro→Ala strains, which correlates with the impact each mistranslating tRNA has on growth. In contrast to the upregulation of proteasome components, proteins involved with autophagy were not induced in response to Pro→Ala or Arg→Ser mistranslating tRNA variants and only four autophagy-related proteins were very weakly up-regulated in the Pro→Ser strain (Figure S2B) suggesting that the mis-made proteins created by mistranslating tRNAs are largely dealt with by the proteasome. Overall, these results demonstrate that mistranslation leads to upregulation of protein chaperones and the proteasome, consistent with previous transcriptomic studies of the mistranslation response (Paredes *et al*. 2012; Reverendo *et al*. 2014; Berg *et al*. 2021b; Hou *et al*. 2023). Furthermore, our results demonstrate that the magnitude of this response at the proteome level differs depending both on mistranslation frequency and the type of substitution created by the tRNA variant.

### Phosphoproteome changes in response to mistranslating tRNA variants uncover cell cycle dysregulation

Next, we investigated how phospho-signaling pathways are modulated in response to mistranslation by identifying phosphosite abundance changes in each mistranslating tRNA strain relative to the empty vector control strain. Phosphosite abundance was corrected based on protein abundance to accurately capture changes in protein phosphorylation, rather than simply changes in protein abundance. Phosphosite abundances can be found in Table S5 and the fold-change and adjusted *p*-values for each phosphosite in each condition can be found in Table S6. Similar to the changes observed at the proteome level, the Pro→Ser strain had the most differentially regulated phosphosites (|fold-change| > 1.5, adjusted *p*-value < 0.05) after controlling for protein abundance changes with 405 sites increasing and 253 sites decreasing. The Pro→Ala and Arg→Ser strain both had fewer differentially regulated phosphosites, with 92 and 53 sites changing in abundances, respectively. Clustering the phosphosite responses, we observed that the sites regulated in both the Pro→Ala and Arg→Ser strains tended to be regulated in the same direction as in the Pro→Ser strain (Figure 4A), suggesting that the phospho-signaling response to mistranslating tRNA variants is largely similar but differs in magnitude between variants. While gene ontology enrichment analysis of the clusters yielded relatively few results, down-regulated phosphosites were significantly enriched on proteins involved in cytokinesis and budding, suggesting mistranslation may impact cell cycle.

**Figure 4.**
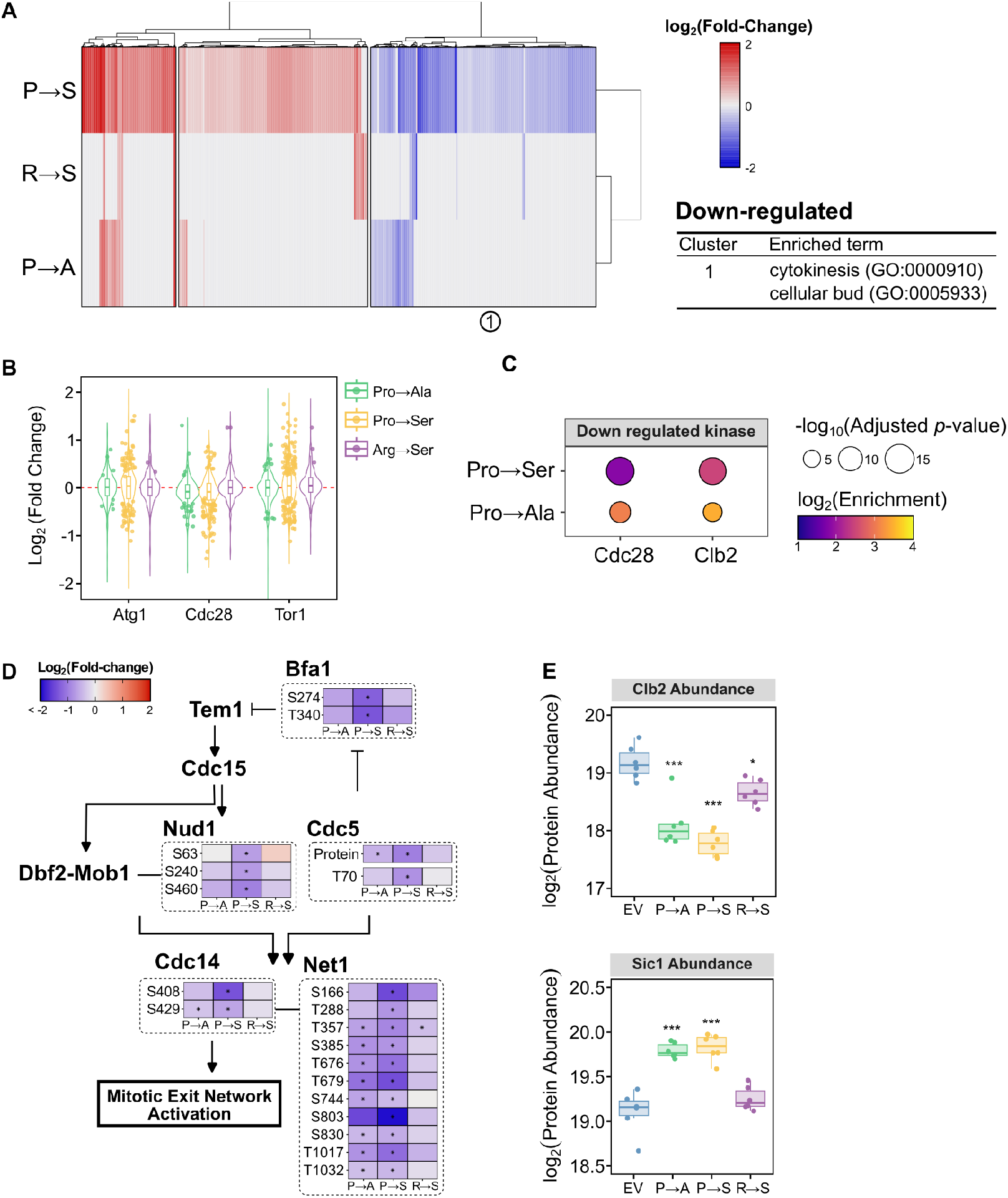
Phosphorylation changes in response to mistranslation uncover cell cycle dysregulation. **(A)** Heatmap of hierarchically clustered differentially abundant phosphosites (adjusted *p*-value < 0.05 in at least one strain) for each mistranslating strain relative to the control. Fold-change for each gene is the average of six replicates. Up-regulated phosphosites are colored in red while down-regulated phosphosites are colored blue. Significantly enriched GO terms relative to all quantified phosphoproteins was determined using g:Profiler (Kolberg *et al*. 2023). **(B)** Violin plot representing the fold change of all measured phosphosites regulated by specific kinases in each mistranslating strain relative to the empty vector control strain. Points show significantly different phosphosites (adjusted *p*-value < 0.05). **(C)** Kinase-substrate enrichment analysis of up and down-regulated phosphosites. Only statistically significantly enriched kinases (adjusted *p*-value < 0.05) with more than 4 regulated phosphosites are shown. **(D)** Regulated phosphosites on proteins involved in the mitotic exit network. Statistically significant changes in phosphosite abundance relative to an empty vector control, after correcting for protein abundance, are marked with a star (adjusted *p*-value < 0.05). **(E)** Boxplot showing protein abundance changes for cell cyclin Clb2p and cyclin-dependent kinase inhibitor Sic1. Points represent independent biological replicates (n = 6) and stars represent statistical significance (* adjusted *p* < 0.05, *** adjusted *p* < 0.0005).

To understand the kinases involved in the response to mistranslating tRNAs, phosphosites were grouped by kinase and the fold change of each site relative to the control strain was plotted (Figure S3). In response to all mistranslating tRNAs, many phosphosites targeted by Atg1, Cdc28, and Tor1 both increased and decreased in abundance (Figure 4B). Kinase-substrate enrichment analysis revealed that in both the Pro→Ser and Pro→Ala strains, down-regulated phosphosites were significantly enriched for targets of the main cell cyclin-dependent kinase Cdc28 and the cyclin Clb2 (Figure 4C). Clb2 is a B-type cyclin that accumulates during G2 and activates Cdc28 to promote transition to M phase (Ghiara *et al*. 1991; Surana *et al*. 1991; Fitch *et al*. 1992). Interestingly, we did not observe this enrichment in the Arg→Ser strain. This might suggest that the response is specific to mistranslation at proline codons, or can simply reflect the milder impact of the Arg→Ser tRNA on the proteome and phosphoproteome.

Closer examination of the downregulated phosphosites suggests that Pro→Ser mistranslation, and to a lesser extent Pro→Ala, specifically disrupt mitotic exit network (MEN) activation as seen through changes in phosphorylation of many proteins in the pathway (Figure 4D). In strains experiencing Pro→Ser mistranslation, there is a decrease in abundance of Polo-like kinase Cdc5 and in phosphorylation at T70, which is required for MEN activation (Rodriguez-Rodriguez *et al*. 2016). Downstream of Cdc5, Pro→Ser strains exhibit decreased phosphorylation of sites on Bfa1, Cfi1/Net1 and Cdc14. Bfa1 is a GTPase activating protein that inactivates Tem1, a GTPase that initiates the MEN (Bardin *et al*. 2000). Hu *et al*. (2001) demonstrated that Bfa1 phosphorylation by Cdc5 inhibits Bfa1 activity to promote mitotic exit. Further supporting this idea, we observe decreased phosphorylation of Nud1, a key scaffold protein required for the activation of Dbf2-Mob1 kinase by Cdc15 (Rock *et al*. 2013), in the Pro→Ser strain. This likely also contributes to the decreased phosphorylation of Cfi1/Net1 and Cdc14, a phosphatase that inactivates mitotic CDKs returning the cell to G1 (Shou *et al*. 1999; Visintin *et al*. 1999). At the level of protein abundance, Clb2 levels decrease and levels of the mitotic CDK inhibitor Sic1 increase in the Pro→Ser and Pro→Ala strains (Figure 4E).

The disruption of cell cycle signaling detected in response to Pro→Ser mistranslation, particularly around late anaphase, is consistent with the negative genetic interactions we previously identified with Pro→Ser mistranslation. For example, the growth impact of the Pro→Ser tRNA is exacerbated in genetic backgrounds with impaired chromosome segregation/cytokinesis (*dam1, esp1, cdc11, stu2, spc24, ctf8*, and *sfi1*), defective anaphase-promoting complex (*cdc23* and *cdc20*) and in strains disrupted for main phosphatase involved in the MEN, *cdc14* (Berg *et al*. 2022). These negative genetic interactions are also seen, to a lesser extent, in strains expressing the Pro→Ala tRNA and in strains exposed to the proline analog azetidine-2-carboxylic acid (Berg *et al*. 2020).

Overall, these results demonstrate that mistranslation of either serine or, to a lesser extent, alanine at proline codons lead to changes in phosphorylation on proteins involved in cell cycle. Previous studies have shown high levels of mistranslation at proline codons lead to G1 arrest (Trotter *et al*. 2001; Berg *et al*. 2021a). Here with lower mistranslation levels, we observe reduced phosphorylation specifically associated with MEN. Because these measurements were taken from asynchronous cells, this likely reflects a smaller population of cells at this phase of the cell cycle compared to the control strain.

These results may also explain the partial slow growth phenotype these strains experience.

### Phosphorylation of translation factors in response to mistranslation

In all the mistranslating strains, the most up-regulated phosphosites occurred on elongation factor 2 (eEF2; Eft1; Figure 5A). Specifically, residues S557, S569, S572 and T763 were the most highly phosphorylated in response to all mistranslating tRNA variants. eEF2 is an evolutionary conserved protein that associates with ribosomes to promote GTP-dependent translocation along the mRNA (Moldave 1985). Previous studies in yeast have shown eEF2 is phosphorylated at T57 by Rck2, a Ser/Thr kinase homologous to the mammalian calmodulin kinases (Nairn and Palfrey 1987; Teige *et* al. 2001). This phosphorylation reduces affinity for GTP and decreases ribosome binding (Dumont-Miscopein *et al*. 1994). While the peptide containing T57 was not measured, the mistranslating tRNAs do not activate Rck2, which is mediated through its phosphorylation, suggesting an alternative kinase may be responsible for eEF2 C-terminal phosphorylation.

**Figure 5.**
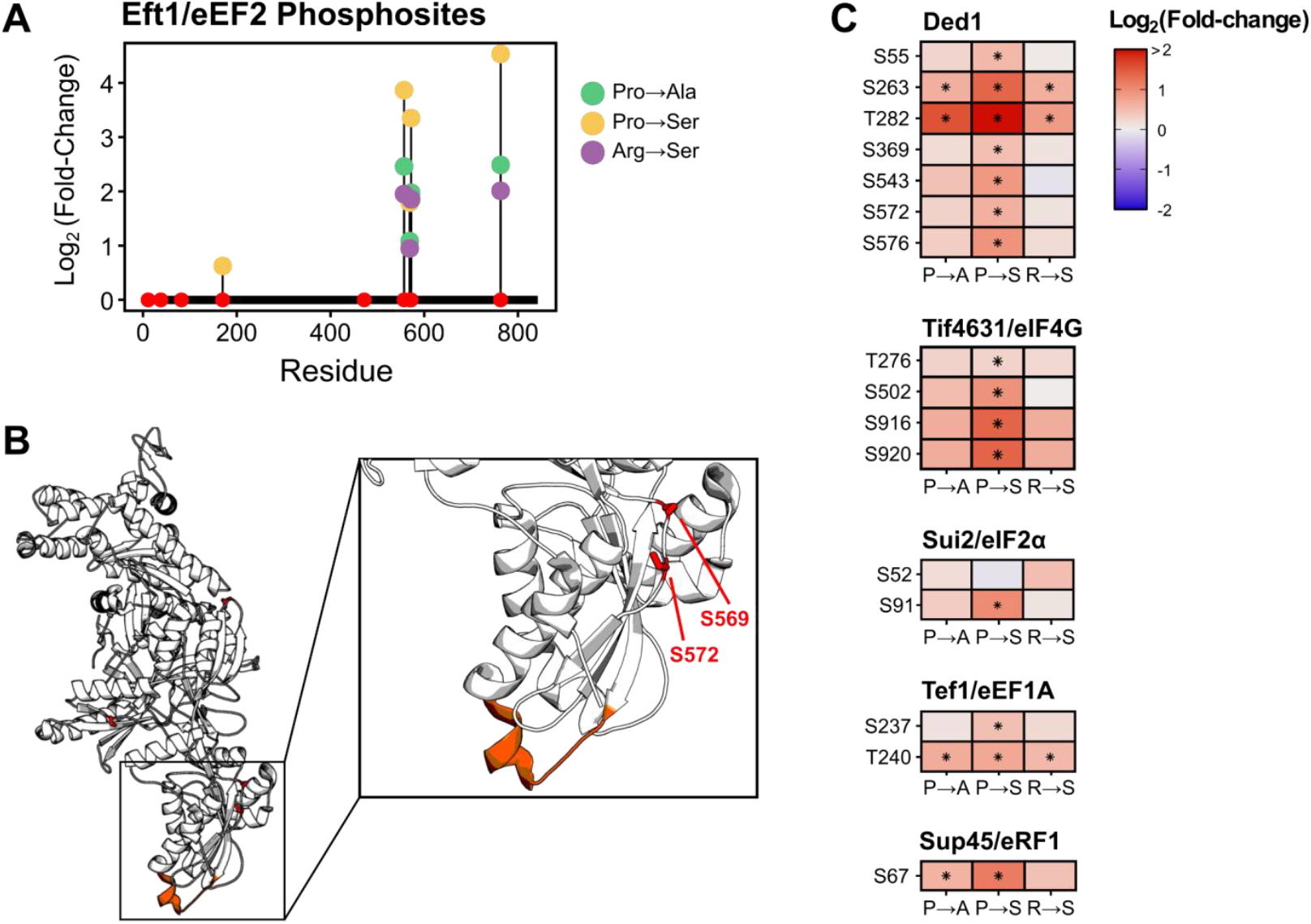
Mistranslation leads to phosphorylation of translation factors. **(A)** Lollipop plot representing phosphorylation sites detected in Eft1. Red points represent all observed phosphosites. Lollipops denote the log_2_ fold change of statistically significant (adjusted *p-*value < 0.05) phosphosites relative to an empty vector control after correcting for protein abundance for the Pro→Ala (green), Pro→Ser (yellow) and Arg→Ser (purple) tRNA variants. **(B)** Structure of Eft1 (PDB: 1N0V; Jørgensen *et al*. 2003) with serine residues that were phosphorylated in the mistranslating strains highlighted in red. Orange region denotes the anticodon mimicry region of domain IV. **(C)** Regulated phosphosites on proteins involved in translation. Statistically significant changes in phosphosite abundance relative to an empty vector control, after correcting for protein abundance, are marked with a star (adjusted *p*-value < 0.05).

Interestingly, two of the up-regulated phosphosites, S569 and S572, occur in domain IV proximal to where eEF2 interacts with the tRNA in the ribosome P-site (Spahn *et al*. 2004; Figure 5B). Ortiz *et al*. (2006) found that mutations in this domain result in sensitivity to translation inhibitors, reduce translation rates and increase -1 frameshifts. The other up-regulated phosphosite, S557, occurs near the sordarin binding region of eEF2 (Jørgensen *et al*. 2003; Søe *et al*. 2007). Sordarin is a tetracyclic diterpene glycoside inhibitor of fungal protein synthesis that bind eEF2 and induce a conformational change to prevent release of eEF2 from the ribosome after translocation (Justice *et al*. 1998; Jørgensen *et al*. 2003; Spahn *et al*. 2004). It is possible that these up-regulated phosphosites modulate translation in response to mistranslation. In agreement with this, expression of mistranslating tRNA in mammalian cells decreases translation rates (Geslain *et al*. 2010; Varanda *et* al. 2020; Lant *et al*. 2021; Hasan *et al*. 2023; Davey-Young *et al*. 2024; Tennakoon *et al*. 2025). In some of these cases, decreased translation rate correlated with eIF2α-S52 phosphorylation but this was not the case for the mistranslating tRNA variants studied here (Figure 5C). Therefore, it is possible C-terminal eEF2 phosphorylation regulates translation in response to mistranslation.

In addition to upregulating eEF2 phosphorylation sites, mistranslation also induced phosphorylation on other translation factors (Figure 5C). These include translation initiation factors Ded1, an ATP-dependent DEAD-box helicase which promotes translation pre-initiation complex assembly (Chuang *et al*. 1997; de la Cruz *et al*. 1997) and Tif4631/eIF4G, a subunit of the eIF4F cap-binding protein complex that works with Ded1 to accelerate assembly of the 48S preinitiation complex (Goyer *et al*. 1993; Gupta *et al*. 2018). There is also increased phosphorylation on the translation elongation factor EF-1α protein Tef1 and the eRF1 polypeptide release factor Sup45. While we do not observe upregulation of eIF2α (Sui2) S52, we do detect increased phosphorylation on S91.

Overall, these results suggest that mistranslating tRNAs could impact translation through increased phosphorylation of translation factors. However, additional experiments are required to determine how phosphorylation at these sites modulates factors like translation rates, processivity and fidelity as well as the upstream kinases responsible for phosphorylating these targets.

### Mistranslating serine tRNA variants create novel phosphorylation sites

Both the Pro→Ser and Arg→Ser tRNAs mis-insert serine into proteins, creating new phospho-acceptor sites. To determine if these positions are phosphorylated, we searched the DDA phosphopeptide enriched samples with phosphorylation, mistranslation and the combined mistranslated-phosphorylation as variable modifications. We stringently filtered the data to only retain peptide sequences that were observed in all six replicates. Indeed, we observed 128 phosphorylation events on mis-inserted serine residues in the Pro→Ser strain and 16 in the Arg→Ser strain (Figure 6A; Table S7). Example spectra of representative novel phosphopeptides are shown in Figure 6B, C. The new phosphosites tended to be in exposed protein regions and had similar solvent accessibility compared to canonical phosphosites detected on STY residues (Figure S4A). This result was not simply due to bias in the mistranslated positions observed in the global proteome data, as we detected mistranslation at proline and arginine residues that spanned the entire range of solvent accessibility (Figure S4B). Additionally, the new phosphosites tend to be on slightly more abundant proteins compared to all detected phosphoproteins (Figure S4C), consistent with mistranslation being more frequently detected in abundant proteins (Figure S4D). Motif enrichment revealed that new phosphosites created by Pro→Ser mistranslation were statistically enriched at PX[P→pS] and [P→pS]P motifs (*p*-value < 10^−6^; Figure 6D). The proline directed SP motif is characteristic of the CMGC family of kinases, which includes cyclin-dependent kinases (CDKs), CDK-like kinases, and mitogen-activated protein kinases (Pinna and Ruzzene 1996). Interestingly, while no significant motif was enriched for the new phosphosites created by Arg→Ser mistranslation, 10 of the 16 novel phosphosites observed in all six biological replicates were followed by a proline residue (i.e. [R→pS]P motif).

**Figure 6.**
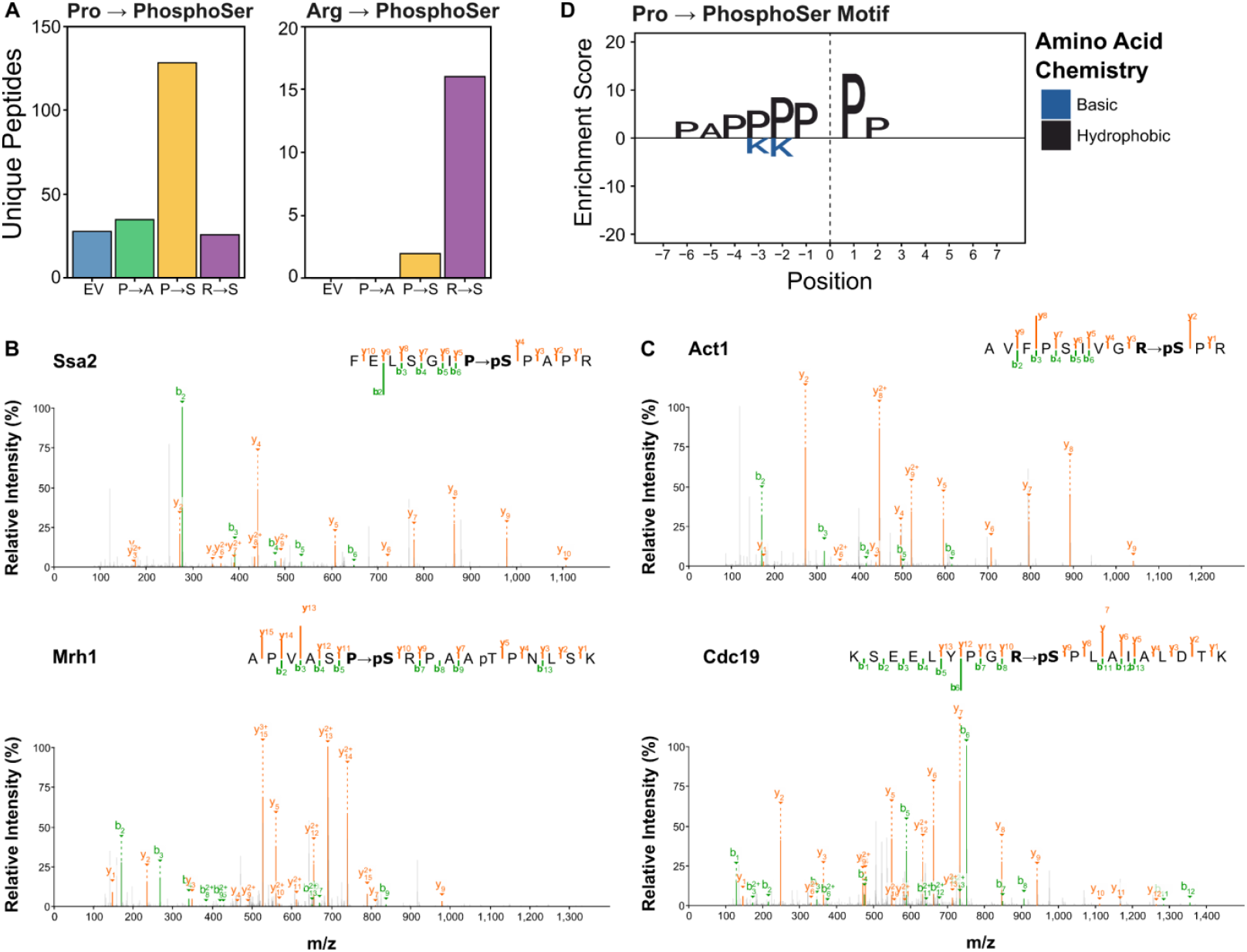
Mistranslating serine at proline or arginine codons creates new phosphoserine sites. **(A)** Bar plot representing the total number of unique peptides with either proline (left) or arginine (right) to phosphoserine modifications. Modified peptides must have been observed in all six biological replicates to be counted. **(B)** Representative spectra for peptides with proline to phosphoserine modifications. Top spectrum shows Ssa2 peptide with proline to phosphoserine modification at residue 462 and bottom spectrum shows Mrh1 peptide with proline to phosphoserine modification at residue 290. **(C)** Representative spectra for peptides with arginine to phosphoserine modifications. Top spectrum shows Act1 peptide with arginine to phosphoserine modification at residue 37 and bottom spectrum shows Cdc19 peptide with arginine to phosphoserine modification at residue 77. **(D)** Enriched amino acids around the proline to phosphoserine modified residue.

Based on these results, it is possible that in addition to directly altering protein sequence, mistranslation can also lead to novel posttranslational modification sites that could impact protein function. For example, we detect a new phosphosite on Protein Kinase C (Pkc1) in the region that interacts with activator Rho1 (Nonaka *et al*. 1995), which could modulate Pkc1 activity and alter downstream signaling. Additionally, a new phosphosite at position 179 in Sir3, a component of the chromatin silence complex, occurs in the bromoadjacent homology domain responsible for nucleosome binding (Onishi et al. 2007). Mutation of this position to leucine acts dominantly to disrupt DNA silencing (Buchberger *et al*. 2008). Therefore, it is possible that creation of a new phosphosite acts similarly.

We also identify new sites on proteins involved in processes impacted by the mistranslating tRNAs. These include sites on chaperones/co-chaperones (Ssa2, Hsp42, Hsc82, Zuo1, Ydj1), translation initiation (Fun12) and elongation factors (Tif35, Tif3, Tef4), ribosomal proteins (Rpp0, Rpp2a, Rpp2b, Rps3, Rps9b, Rps10a, Rps16a, Rpl3, Rpl24b, Rpl32), and a protein involved in regulating translation (Stm1). Overall, our results demonstrate that mistranslation can introduce new phospho-acceptor sites that become phosphorylated with the potential to influence protein function.

## CONCLUSIONS

Overall, our results demonstrate that mistranslating tRNA variants impact the abundance of ∼300 proteins and phosphorylation of ∼450 proteins at the global level. The proteins and phosphosites that change in response to mistranslation tend to be similar between different mistranslating tRNA variants. By comparing different mistranslating tRNAs, we find the number of differentially abundant proteins and phosphosites correlate with the impact of each tRNA variant on growth. Both growth impact and magnitude of proteome/phosphoproteome changes differ depending on the type of substitution created. Our findings demonstrate that mistranslating tRNAs alter the abundance and phosphorylation of proteins involved in key cellular processes, including proteostasis, cell cycle, and translation. Modulation of these processes must be considered when investigating the contribution of naturally occurring mistranslating tRNA variants to disease (Berg *et al*. 2019a; Tennakoon *et al*. 2025) and when developing mistranslating tRNAs to be used as therapeutics (Lueck *et al*. 2019; Hou *et al*. 2023; Porter *et al*. 2024; Beharry *et al*. 2025).

## DATA AVAILABILITY

Strains and plasmids are available upon request. The authors affirm that all data necessary for confirming the conclusions of the article are present within the article, figures, and supplemental material. Supplementary File 1 contains all supplemental figures. Supplemental File 2 contains all the supplemental tables with mistranslation frequencies, protein/phosphopeptide quantification, fold-change values and p-values. The mass spectrometry proteomics data have been deposited to the ProteomeXchange Consortium via the PRIDE (Perez-Riverol *et al*. 2019) partner repository with the dataset identifier PXD068388 and PXD068392. An annotated list of all the raw files can be found in Supplemental File 3. Report logs containing full parameters for the raw data analysis, DIA libraries, the DIA-NN/MSFragger outputs and custom R scripts used to analyze the data and create the figures can be found at https://github.com/Villen-Lab/Mistranslation-Phospho-Proteome-AnalysisFiles.

## ACKNOWLEDGMENTS

We thank Ariadna Llovet and Julian Ramos for providing support with in-house proteomics software and Alex Hogrebe for instrumentation illustrations. We would also like to thank Chris Brandl, Josh Isaacson and Ecaterina Cozma for critically reading the manuscript.

