## Supplemental File 1 for "Mistranslating tRNA variants impact the proteome and phosphoproteome of *Saccharomyces cerevisiae*"

### Supplemental Figures

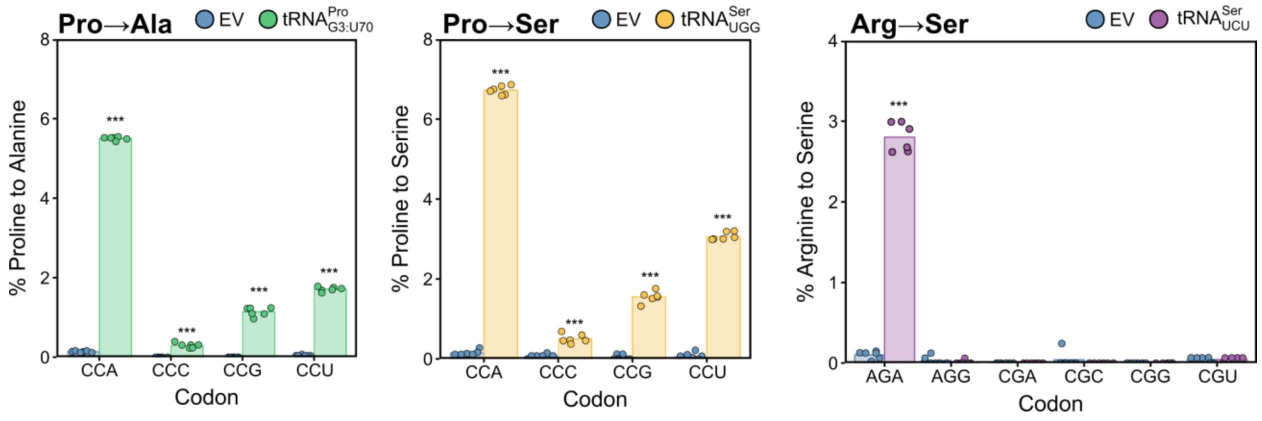


**Figure S1. Codon specific mistranslation frequency.** For each mistranslating tRNA variant, the number of mistranslation events at each synonymous codon was determined relative to the total number of wild-type residues observed per codon. Each point represents one biological replicate. Strains expressing a mistranslating tRNA were compared to the empty vector control strain (EV) using a *t*-test with Bonferroni correction. * $P < 0.01$; ** $P < 0.001$; *** $P < 0.0001$.

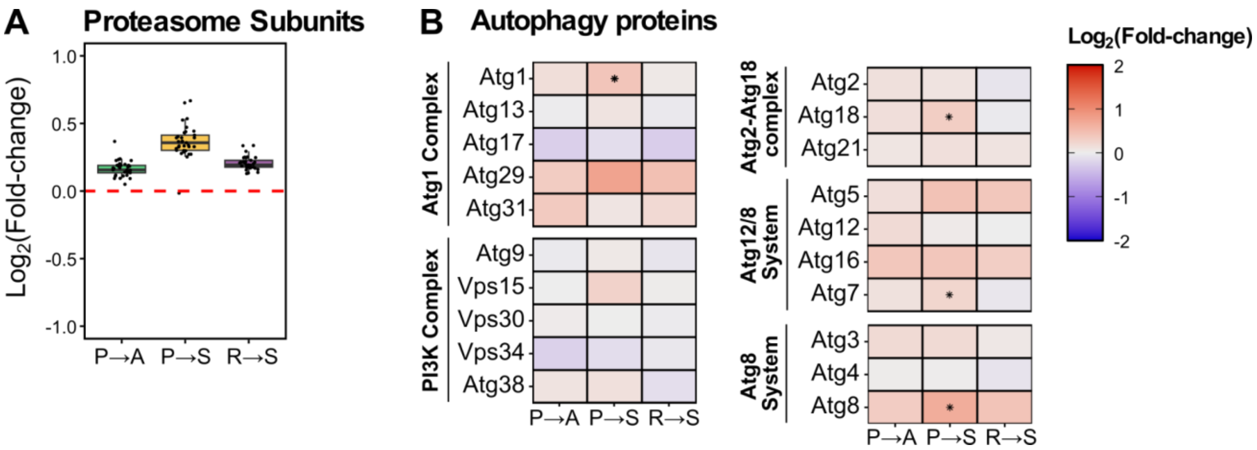


**Figure S2. Impact of mistranslating tRNA variants on proteasome and autophagy-related protein abundance. (A)** Box plot showing the $log_2$-fold change in protein abundance for proteasome subunits relative to an empty vector control strain. Each point represents a single proteasomal protein. The dotted red line indicates no change relative to the empty vector control strain. **(B)** Heat maps representing the $log_2$-fold change in protein abundance for selected proteins involved in autophagy. Stars indicate statistically significant changes in protein abundance relative to an empty vector control strain (adjusted *p*-value < 0.01).

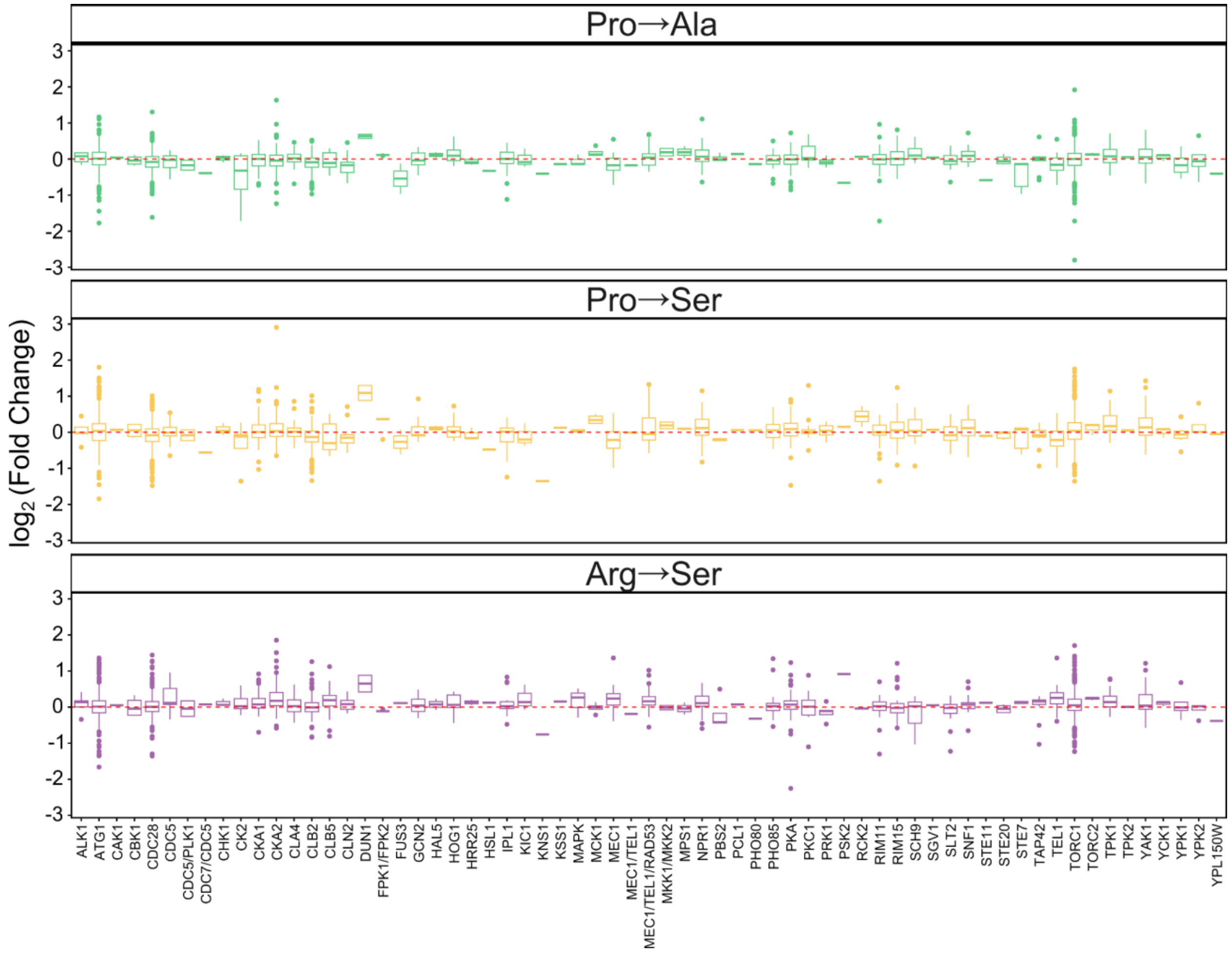


**Figure S3. Fold change of phosphosites grouped by kinase.** Box plots representing the fold change of all measured phosphosites regulated by specific kinases in each mistranslating strain relative to the empty vector control strain. The dotted line represents zero fold change.

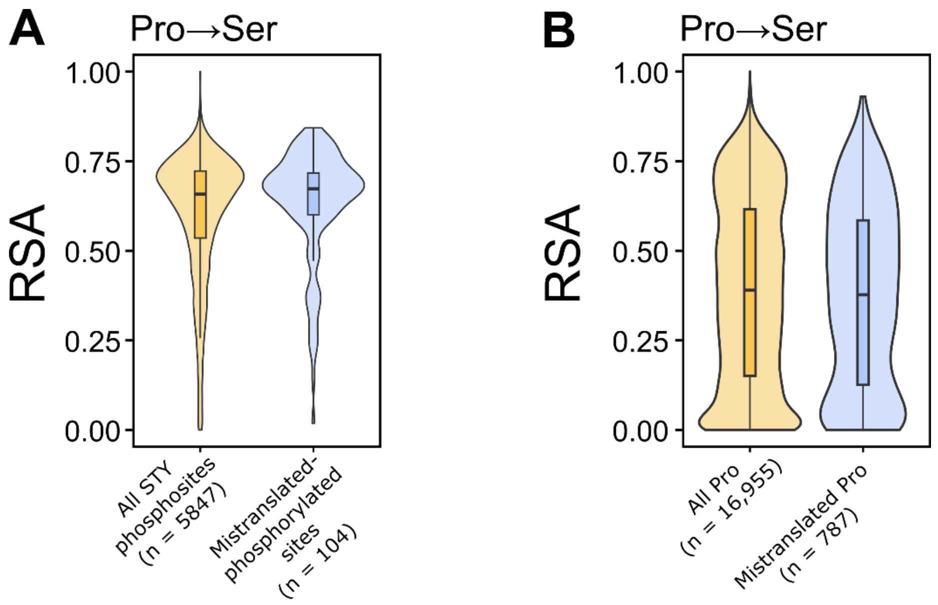


**Figure S4. New phosphosites created by mistranslation have similar accessibility to canonical phosphosites on STY residues. (A)** Violin plot representing the relative solvent accessibility (RSA) of all measured phosphorylated STY residues and new phosphorylated sites at mistranslated positions. Relative solvent accessibility was determined from AlphaFold2 structural models. To be included, peptides had to be present in all six biological replicates. **(B)** Violin plot representing the RSA of all measured proline residues as well as mistranslated residues. Relative solvent accessibility was determined from AlphaFold2 structural models. To be included, peptides had to be present in all six biological replicates.

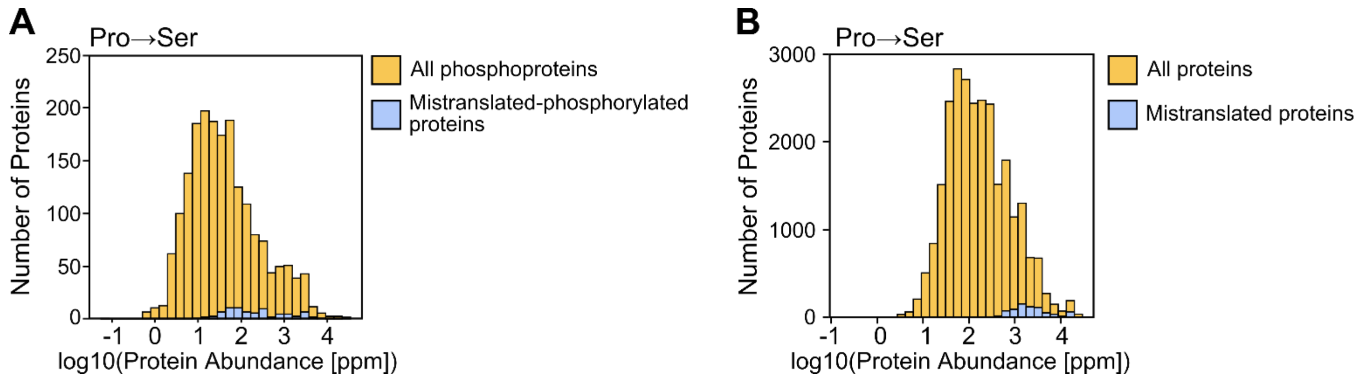


**Figure S5. New phosphosites created by mistranslation are detected more frequently on higher abundance proteins. (A)** Histogram representing the abundance of phosphoproteins and proteins with new phosphorylation sites created by mistranslation. **(B)** Histogram representing the abundance of all detected proteins and proteins where we detect mistranslation of Pro→Ser. Protein abundance values were obtained from the Protein Abundance Database (https://pax-db.org/).
